# Chemical-genetic interactions with the proline analog L-azetidine-2-carboxylic acid in *Saccharomyces cerevisiae*

**DOI:** 10.1101/2020.08.10.245191

**Authors:** Matthew D. Berg, Yanrui Zhu, Joshua Isaacson, Julie Genereaux, Raphaël Loll-Krippleber, Grant W. Brown, Christopher J. Brandl

## Abstract

Non-proteinogenic amino acids, such as the proline analog L-azetidine-2-carboxylic acid (AZC), are detrimental to cells because they are mis-incorporated into proteins and lead to proteotoxic stress. Our goal was to identify genes that show chemical-genetic interactions with AZC in *Saccharomyces cerevisiae* and thus also potentially define the pathways cells use to cope with amino acid mis-incorporation. Screening the yeast deletion and temperature sensitive collections, we found 72 alleles with negative synthetic interactions with AZC treatment and 12 alleles that suppress AZC toxicity. Many of the genes with negative synthetic interactions are involved in protein quality control pathways through the proteasome. Genes involved in actin cytoskeleton organization and endocytosis also had negative synthetic interactions with AZC. Related to this, the number of actin patches per cell increases upon AZC treatment. Many of the same cellular processes were identified to have interactions with proteotoxic stress caused by two other amino acid analogs, canavanine and thialysine, or a mistranslating tRNA variant that mis-incorporates serine at proline codons. Alleles that suppressed AZC-induced toxicity functioned through the amino acid sensing TOR pathway or controlled amino acid permeases required for AZC uptake.

## INTRODUCTION

Non-proteinogenic amino acids can be recognized by aminoacyl-tRNA synthetases, charged onto tRNAs and mis-incorporated into proteins [reviewed in Rodgers and Shiozawa (2008)]. L-azetidine-2-carboxylic acid (AZC) is a non-proteinogenic imino acid analog of proline produced by liliaceous plants as well as garden and sugar beets (Fowden 1959, 1972; Rubenstein *et al*. 2006). In species where AZC is not normally produced, it is activated by the prolyl-tRNA synthetase, charged onto tRNA^Pro^ and mis-incorporated into proteins at proline codons (Peterson and Fowden 1963). In nature, AZC inhibits growth of surrounding vegetation and poisons predators presumably by inducing protein mis-folding (Fowden and Richmond 1963). In mammals, AZC treatment leads to defects in the production of the proline rich protein collagen and results in limb deformation (Aydelotte and Kochhar 1972).

The proteotoxic stress that results from incorporating non-proteinogenic amino acids into proteins is similar to that arising from errors in protein synthesis during translation. Mistranslation occurs at a frequency of 10^−4^ to 10^−6^ (Joshi *et al*. 2019), and increases in response to different environmental conditions (Ling and Söll 2010; Wang and Pan 2016; Mohler *et al*. 2017) or due to mutations in the translational machinery (Moghal *et al*. 2014; Raina *et al*. 2014; Hoffman *et al*. 2017; Lant *et al*. 2017; Berg *et al*. 2017). Mis-incorporation frequencies as high as 8 to 10% can be achieved in yeast and *Escherichia coli* (Ruan *et al*. 2008; Mohler *et al*. 2017; Zimmerman *et al*. 2018; Berg *et al*. 2019).

The goal of our screen was to identify genes that show chemical-genetic interactions with AZC in *Saccharomyces cerevisiae* and thus systematically define the potential pathways cells use to cope with amino acid mis-incorporation. Using the yeast deletion (Giaever *et al*. 2002) and temperature sensitive collections (Costanzo *et al*. 2016), we identified 72 alleles with negative synthetic interactions with AZC and 12 alleles that suppressed AZC-induced growth defects. Negative interactions identified the proteasome and genes involved in endocytosis and actin cytoskeletal organization as required for cells to tolerate AZC. These pathways were also required for tolerance to two other non-proteinogenic amino acids, canavanine, an arginine analog, and thialysine, a lysine analog, and to mistranslation resulting from a tRNA^Ser^ variant that mis-incorporates serine at proline codons. Suppressors of AZC-induced toxicity functioned mainly in the TOR signaling pathway and in controlling amino acid permeases.

## MATERIALS AND METHODS

### Genetic screen for AZC interactions

The yeast deletion collection (Giaever *et al*. 2002) and temperature sensitive mutants (Ben-Aroya *et al*. 2008; Li *et al*. 2011; Kofoed *et al*. 2015; Costanzo *et al*. 2016; Berg *et al*. 2018) were arrayed in quadruplicate 1536 format with four replicate colonies for each mutant using a BioMatrix (S&P Robotics Inc.) automated pinning system on yeast extract-peptone medium with 2% glucose (YPD) containing 200 μg/mL geneticin (G418; Invitrogen). Newly pinned arrays were grown for 24 hours then used as the source to re-pin the cells first on synthetic defined (SD) medium containing uracil, leucine, histidine and methionine then on to the same medium containing 30 μg/mL AZC (ChemCruz). Cells were grown for 5 days at 30°. Plates were photographed every 24 hours after pinning.

### Calculating fitness in AZC and identification of hits

Fitness of each strain was defined as the ratio between colony size in medium containing AZC compared to colony size on medium lacking AZC. Raw colony size for each strain was determined using SGATools (Wagih *et al*. 2013). Further data analysis was performed using a custom R script (Supplemental File 1). Strains displaying small colony size in the absence of AZC, defined as growth less than 5% of the average colony size per plate, were removed from the analysis. Fitness of each strain was normalized using a Z-score method per plate. Briefly, the colony sizes were transformed so that the average colony size per plate was 0 and the standard deviation was set to 1. Strains with a Z-score greater than or equal to 3 were called suppressor hits; strains with Z-scores less than or equal to −1.5 were considered to have a negative synthetic interaction. Alleles of essential genes were grouped together and one Z-score calculated for the gene. Data can be found in supplemental file 2.

### Growth curve validation in AZC, canavanine and thialysine-containing media

Negative synthetic and suppressor strains were validated in two independent growth curve experiments. In each experiment, strains were grown in biological triplicate overnight in YPD medium containing 200 μg/mL G418. Cells were washed with sterile water then resuspended in SD medium. Cultures were diluted to OD_600_ of 0.1 in either SD medium or SD medium containing 10 μg/mL AZC. Cells were incubated with agitation at 30° and OD_600_ measured every 15 minutes using a BioTek Epoch 2 microplate spectrophotometer for 24 hours. Area under the curve (AUC) was calculated for each strain using the R package ‘growthcurver’ (Sprouffske and Wagner 2016). The AUC values were used as a measure of fitness and normalized so that the wild-type strain in SD medium had a fitness of 1. The experimental fitness of each mutant grown in AZC medium (W_ij_) was compared to the expected fitness based on the fitness of the wild-type strain grown in AZC medium (W_i_) and the fitness of the mutant strain in SD medium (W_j_) using a multiplicative model (calculated as W_*ij*_ − [W_*i*_ * W_*j*_]). Strains in both replicate experiments that scored less than −0.1 (*p* < 0.05; Welch’s t-test) were considered true negative interactors while strains that scored greater than 0.1 (*p* < 0.05; Welch’s t-test) were considered true positive interactors. Data can be found in supplemental file 3.

Growth curves were performed as above in SD medium or SD medium containing 1.0 μg/mL L-canavanine (Sigma) or 7.5 μg/mL S-aminoethyl-L-cysteine (thialysine; Sigma). Interaction scores for the chemical-genetic interactions were calculated as above. Data can be found in supplemental file 4.

### Synthetic genetic array analysis of AZC hits with mistranslating tRNA^Ser^_UGG, G26A_

The SGA starter strain, Y7092 (*MAT*α *can1*Δ::*STE2pr-SpHIS5 lyp1*Δ *his3*Δ*1 leu2*Δ*0 ura3*Δ*0 met15*Δ*0*), was a kind gift from Dr. Brenda Andrews (University of Toronto). A wild-type tRNA^Ser^ or tRNA^Ser^_UGG, G26A_, a serine tRNA with a proline anticodon and a G26A mutation (Berg *et al*. 2017), were integrated at the *HO* locus with the *natNT2* cassette to create CY8611 (*MAT*α *HO::natNT2 can1*Δ::*STE2pr-SpHIS5 lyp1*Δ) and CY8613 (*MAT*α *HO::natNT2-SUP17_UGG, G26A_ can1*Δ::*STE2pr-SpHIS5 lyp1*Δ) as described in Zhu *et al*. (2020).

The SGA analysis was performed as described by Tong *et al*. (2001). Briefly, strains that showed negative synthetic interactions with AZC were re-arrayed in 384 format using the BioMatrix automated pinning robot. Three arrays were constructed. Each validated AZC sensitive allele was surrounded by a control *his3Δ* strain to mitigate nutrient effects. The position of each AZC hit was randomized on each of the three arrays. The arrays were expanded to 1536-format where each strain was represented in quadruplicate on each plate. The SGA control and query strains (CY8611 and CY8613) were mated to each array. Mated strains were grown overnight then pinned onto YPD plates containing 100 μg/mL nourseothricin-dihydrogen sulfate (clonNAT; Werner BioAgents) and 200 μg/mL G418 to select for diploids. Haploids were generated by pinning the diploid strains onto enriched sporulation plates and incubating for 1 week at 22°. The haploids then underwent three rounds of selection for double mutants that had both the tRNA mutation and temperature-sensitive allele. First, strains were pinned on SD plates lacking histidine, arginine and lysine and containing canavanine and thialysine to select for MAT*a* haploids. Next, colonies were pinned twice onto SD plates containing MSG as the nitrogen source, lacking histidine, arginine and lysine and containing canavanine, thialysine, G418 and clonNAT to select double mutants. Colonies were incubated for two days between pinnings at room temperature. The double mutants were grown at 30° for 5 days and imaged every 24 hrs. Images from day 3 were analyzed and scored using SGAtools (Wagih *et al*. 2013) and a custom R script (supplemental file 1). Alleles with SGA score ≤ −0.1 and a corrected p-value ≤ 0.05 were called as synthetic with tRNA^Ser^_UGG, G26A_. Data can be found in supplemental file 5.

### Fluorescent microscopy and actin patch quantification

A wild-type strain was grown overnight in SD media and diluted to an OD_600_ of 0.1 in either SD media or SD media containing various concentrations of AZC (1 μg/mL, 2.5 μg/mL, 6 μg/mL, 10 μg/mL). Cells were grown to an OD_600_ of 1 before being fixed in 3.7% formaldehyde (Sigma). Cells were stained with fluorescein isothiocyanate-ConA, rhodamine-phalloidin, and 4’,6-diamidino-2-phenylindole as described in Ohya *et al*. (2005) and imaged with 100x magnification on a Zeiss Axio Imager Z1 Fluorescent microscope using ZEN Blue Pro software (Zeiss Inc.). The number of actin patches per cell was measured using CalMorph (v1.2; Ohya *et al*. 2005).

### GO term analysis, genetic interaction profiling and SAFE analysis

GO term analysis was performed using the GO term finder tool (http://go.princeton.edu/) using a P-value cut off of 0.01 after applying Bonferroni correction. Terms were filtered with REVIGO (Supek *et al*. 2011). GeneMANIA (Warde-Farley *et al*. 2010) was used to generate protein-protein and genetic interaction networks using the genetic interaction data from Costanzo *et al*. (2010, 2016) and all available protein-protein interactions (GeneMANIA datasets as of November 2019). Networks were constructed using Cytoscape 3.7 (Shannon *et al*. 2003).

Spatial analysis of functional enrichment (SAFE; Baryshnikova 2016) analysis was performed through TheCellMap (http://thecellmap.org; Usaj *et al*. 2017).

### Data Availability

Strains and plasmids are available upon request. The authors affirm that all data necessary for confirming the conclusions of the article are present within the article, figures, and tables. Supplemental data can be found on Figshare. Supplemental materials document contains all supplemental figures. Supplemental file 1 contains custom R script used to analyze the data. Supplemental file 2 contains the raw Z-scores from the initial AZC screen. Supplemental file 3 contains the AUC values and interaction scores for the AZC growth curve validation. Supplemental file 4 contains the AUC values and interaction scores for the canavanine and thialysine growth curves. Supplemental file 5 contain the results from the synthetic genetic array with AZC sensitive strains and the mistranslating tRNA^Ser^_UGG, G26A_.

## RESULTS AND DISCUSSION

### A screen for genes that have chemical-genetic interactions with the proline analog AZC

To identify genes and pathways that have chemical-genetic interactions with AZC, we measured the effect of AZC on growth of strains in the yeast deletion and temperature sensitive collections, containing 4291 and 1016 alleles respectively. Each strain in the collections was pinned in quadruplicate onto SD medium or SD medium containing 30 μg/mL AZC. We selected 30 μg/mL AZC as it decreased average fitness by ~ 40% as measured by colony size (Figure S1). Fitness for each strain was determined from the ratio of the growth of each colony on medium with AZC to growth on medium without AZC after 5 days at 30°. Fitness values were normalized on a per plate basis and converted to Z-scores (Figure 1A). Nineteen genes (0.3% of the mutants screened) had a Z-score greater than 3, whereas 255 genes (5%) had a score less than −1.5 (Figure 1B; all raw colony sizes and fitness values are found in supplemental file 2). These strains were tested further for their potential suppression of AZC toxicity and for their sensitivity to AZC, respectively.

**Figure 1.**
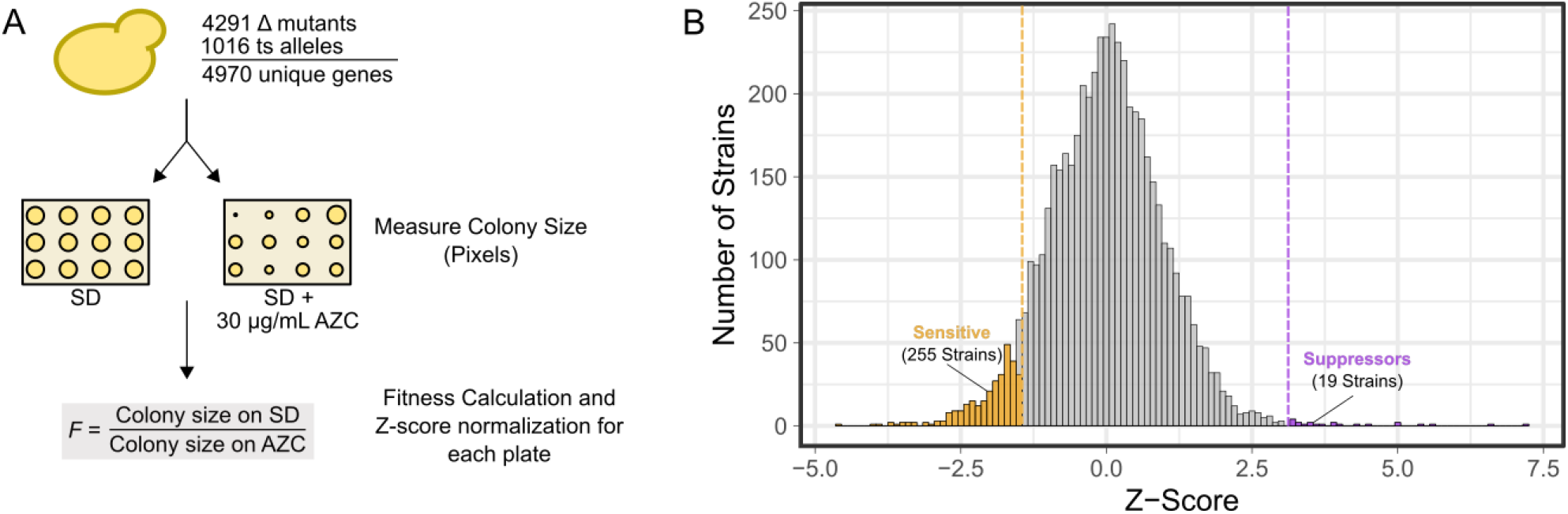
A screen identifying genes that have chemical-genetic interactions with AZC. (A) Overview of the screening approach. (B) Distribution of Z-score normalized fitness values for 5307 strains representing 4970 genes from the deletion and temperature sensitive yeast collections grown on medium containing 30 μg/mL AZC.

The 255 sensitive strains and 19 suppressor strains were analyzed for growth in liquid medium containing AZC. We tested different AZC concentrations with the wild-type strain and a potential AZC sensitive strain *hap5Δ* (Figure S2). The *hap5Δ* strain, with a Z-score of −3.5, was chosen as a representative AZC sensitive strain to ensure that the AZC concentration in the liquid growth assay allowed a dynamic range sufficient to detect differences in sensitivity of the other strains. An AZC concentration of 10 μg/mL was chosen for the validation as it decreased the fitness of the wild-type strain by ~ 40% and the *hap5Δ* strain by greater than 85%. The growth curve analysis in liquid medium confirmed the AZC sensitivity of 72 (28%) strains identified in the screen on solid medium: 43 from the deletion collection and 29 from the temperature sensitive collection (Supplemental File S3). This represents 1.0% and 2.8% of all deletion strains or temperature sensitive strains screened, respectively. The growth curve analysis validated 12 (63%) potential suppressor strains; 6 from the deletion collection and 6 from the temperature sensitive collection.

### Defects in a variety of pathways result in sensitivity to AZC

The gene functions of the 72 AZC sensitive strains fell into 7 main categories when grouped based on their descriptions in the yeast genome database (www.yeastgenome.org; Figure 2A). Genes with a role in protein quality control were the most abundant (17 genes) followed by genes with a role in metabolism/mitochondria (13 genes). GO analysis of biological processes identified an enrichment of genes with roles in the proteasome and actin organization (Figure 2B). Spatial analysis of functional enrichment (SAFE), which detects network areas containing statistically overrepresented functional groups (Baryshnikova 2016), similarly showed enrichment in the protein turnover and cell polarity neighborhood of the yeast genetic interaction network (Figure 2C).

**Figure 2.**
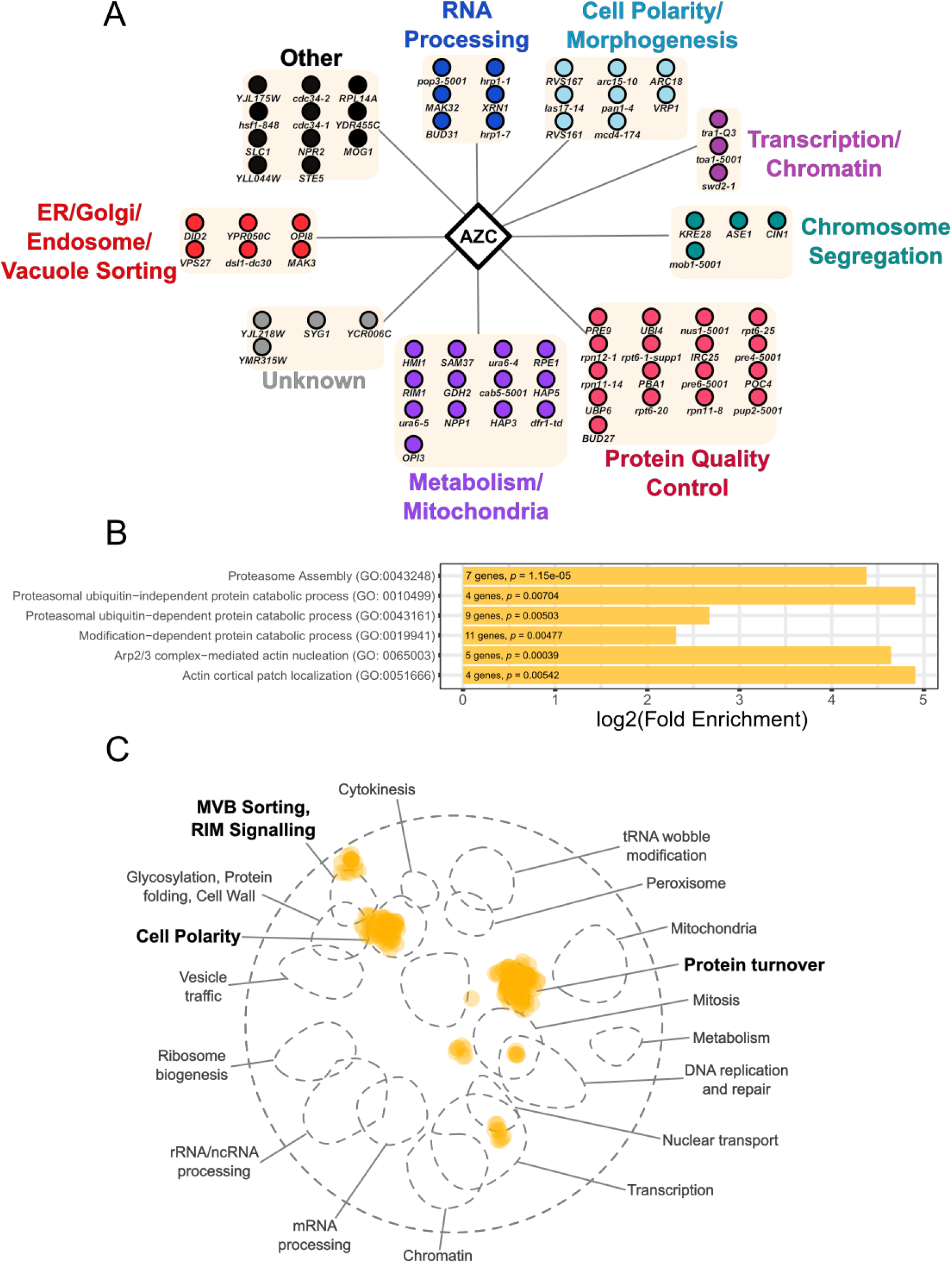
Negative synthetic interactions with AZC. (A) Alleles that have negative synthetic interactions with AZC are arranged according to their predicted function. (B) Significantly enriched GO biological processes were determined from the set of genes with negative synthetic interactions with AZC. The number of genes identified as AZC sensitive corresponding to each GO term and P-values are labelled on each bar. (C) SAFE analysis of genes that have negative synthetic interactions with AZC were mapped onto the yeast genetic interaction profile map (Costanzo *et al*. 2016). Yellow dots represent genes within the local neighborhood of validated AZC sensitive genes. Bold terms represent network regions that are enriched in sensitivity to AZC.

There are multiple mechanisms that could result in a synthetic negative interaction with AZC. First, and most generally, nonredundant genes required for cellular response to proteotoxic stress will be identified since AZC is mis-incorporated into proteins at proline codons. Second, temperature sensitive alleles that are identified could be hypomorphs. Mis-incorporating AZC into the gene product would reduce the steady state level of the protein if the protein had one or more essential proline residues. Related to this, misincorporation into a hypomorphic protein could reduce function sufficiently to cause synthetic interactions with genes (either knockouts or temperature sensitive alleles) in secondary pathways. Third, AZC might act as a chemical inhibitor, independent of its misincorporation into proteins, specifically inhibiting a gene product. This would be similar, for example, to the inhibitory effect of aminotriazole on His3 (Klopotowski and Wiater 1965) and would result in a genetic interaction pattern closely resembling the gene whose protein is the target of AZC.

### Similar cellular processes are required for resistance to other amino acid analogs and tRNA-induced mistranslation

To evaluate if the negative genetic interactions identified here are specific for AZC, we determined the chemical genetic interactions of the 72 AZC-sensitive alleles identified above with canavanine and thialysine, analogs of arginine and lysine, respectively, in liquid media. Both non-proteinogenic amino acids would give rise to proteotoxic stress, but their impact may vary depending upon the number and importance of arginine and lysine residues in specific proteins. From analyzing dose-response curves for each non-proteinogenic amino acid (Figure S3), we selected concentrations of 7.5 μg/mL for thialysine and 1 μg/mL for canavanine. These concentrations resulted in ~ 40% decrease in wild-type growth, similar to the concentration used for AZC. Of the 72 alleles identified to have negative interactions with AZC, 29 were sensitive to thialysine and 19 were sensitive to canavanine (Figure 3; See supplemental file S4 for interaction scores). Fifteen genes were sensitive to all three amino acid analogs.

**Figure 3.**
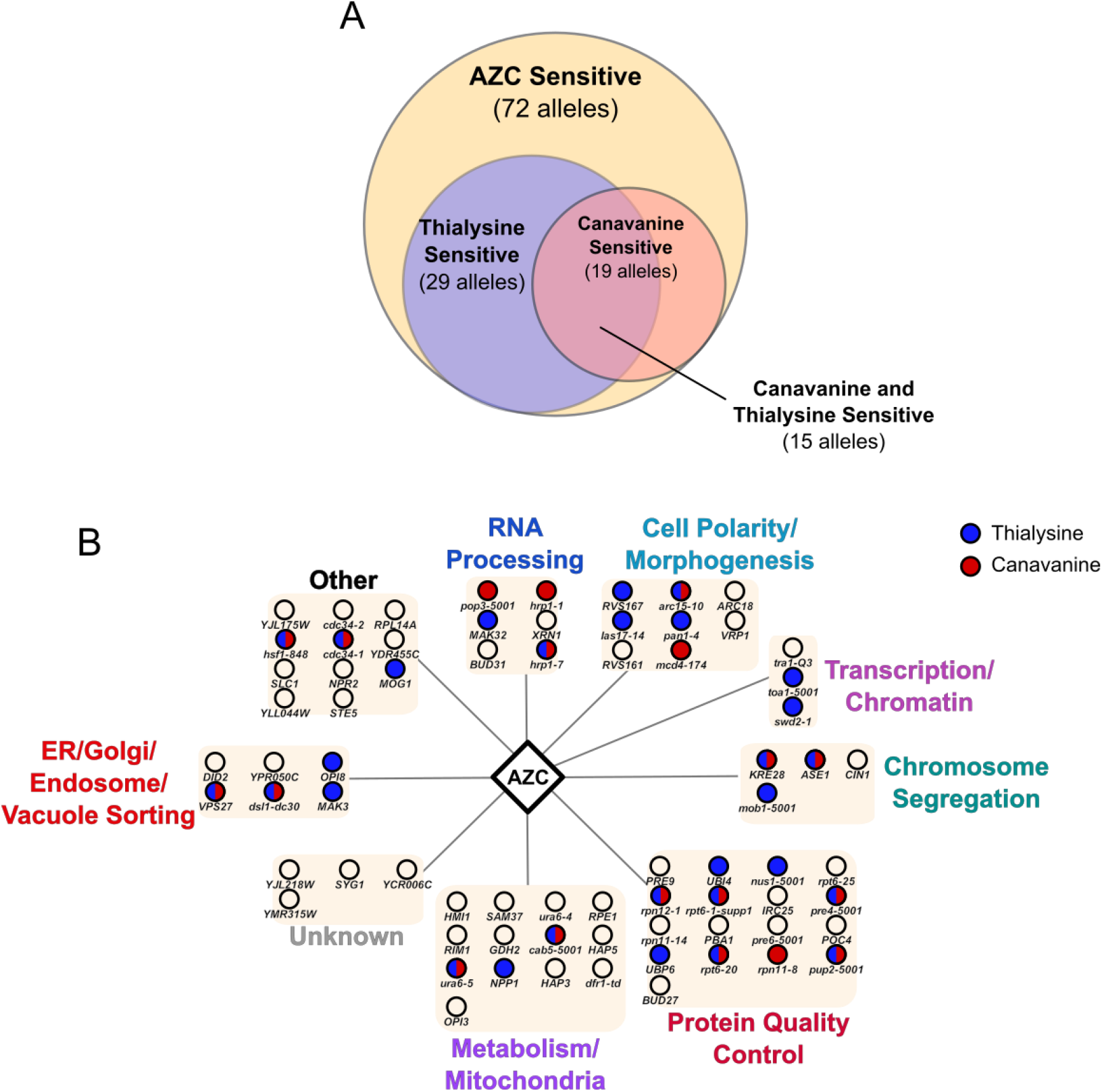
Negative interactions of AZC sensitive genes with thialysine and canavanine. (A) Venn diagram showing the overlap of negative synthetic interactions between AZC, canavanine and thialysine. (B) Alleles that were identified as having negative synthetic interactions with AZC were tested for their sensitivity to thialysine (blue) and canavanine (red) in liquid growth assays.

There were negative synthetic interactions with canavanine and thialysine for genes in all of the functional groups with known functions. For thialysine, the greatest similarity with AZC was seen for genes with roles in ER/golgi/endosome/vacuole sorting (67% overlap), transcription/chromatin (67% overlap), cell polarity/morphogenesis (50% overlap). For canavanine, the greatest similarity with AZC was seen for genes with roles in RNA processing (50% overlap). Both thialysine and canavanine had negative synthetic interactions with genes involved in protein quality control (47% overlap for thialysine and 35% overlap for canavanine). The sensitivity of genes in all these pathways suggests that some genes in these pathways are required to respond to general amino acid mis-incorporation. We do note that although the pathways are similar for the three amino acid analogs, the specific genes identified were not identical. This could suggest that AZC mis-incorporation specifically and uniquely effects some genes in the identified pathways. Whether this represents absolute differences between the effects of AZC and other non-proteinogenic amino acids or differences in dose-response is unclear.

Next, we determined if alleles that have negative interactions with AZC mis-incorporation for proline were also sensitive to mis-incorporation of a canonical amino acid in place of proline. This would provide evidence that native mistranslation provokes similar sensitivities and requires a response similar to the incorporation of a non-proteinogenic amino acid. Using a synthetic genetic array (Tong *et al*. 2001), we measured the fitness of double mutants containing one of the 72 alleles identified above in combination with a mistranslating tRNA^Ser^ variant with a UGG anticodon (Berg *et al*. 2017, 2019). This variant mis-incorporates serine at proline codons at ~ 5% frequency and results in a 20% decrease in growth compared to a wild-type strain (Berg *et al*. 2019). Twenty of the 72 alleles tested had negative genetic interactions with the mistranslating tRNA (Figure 4; See supplemental file S5 for interaction scores). The greatest overlap was seen for alleles with roles in cell polarity and morphogenesis (63%) and, as seen for canavanine and thialysine, protein quality control (41%). Genes in metabolism/mitochondria and transcription/chromatin did not have negative genetic interactions with tRNA derived mistranslation. It is possible the chemical-genetic interactions seen between the metabolism genes and the non-proteinogenic amino acids arise from nutrient signalling effects in response to these supplemented amino acids. It is also possible that tRNA^Ser^_UGG, G26A_ is not imported into the mitochondria, as there are two mitochondrial encoded tRNA^Ser^ genes in yeast (Turk *et al*. 2013). Whether the differences between tRNA derived mistranslation and AZC for genes in the other functional groups arise because of different impacts of AZC and serine on proteins or differences in dose-response is unclear, yet the overlap with AZC sensitive strains is evidence that many of the major pathways are shared.

**Figure 4.**
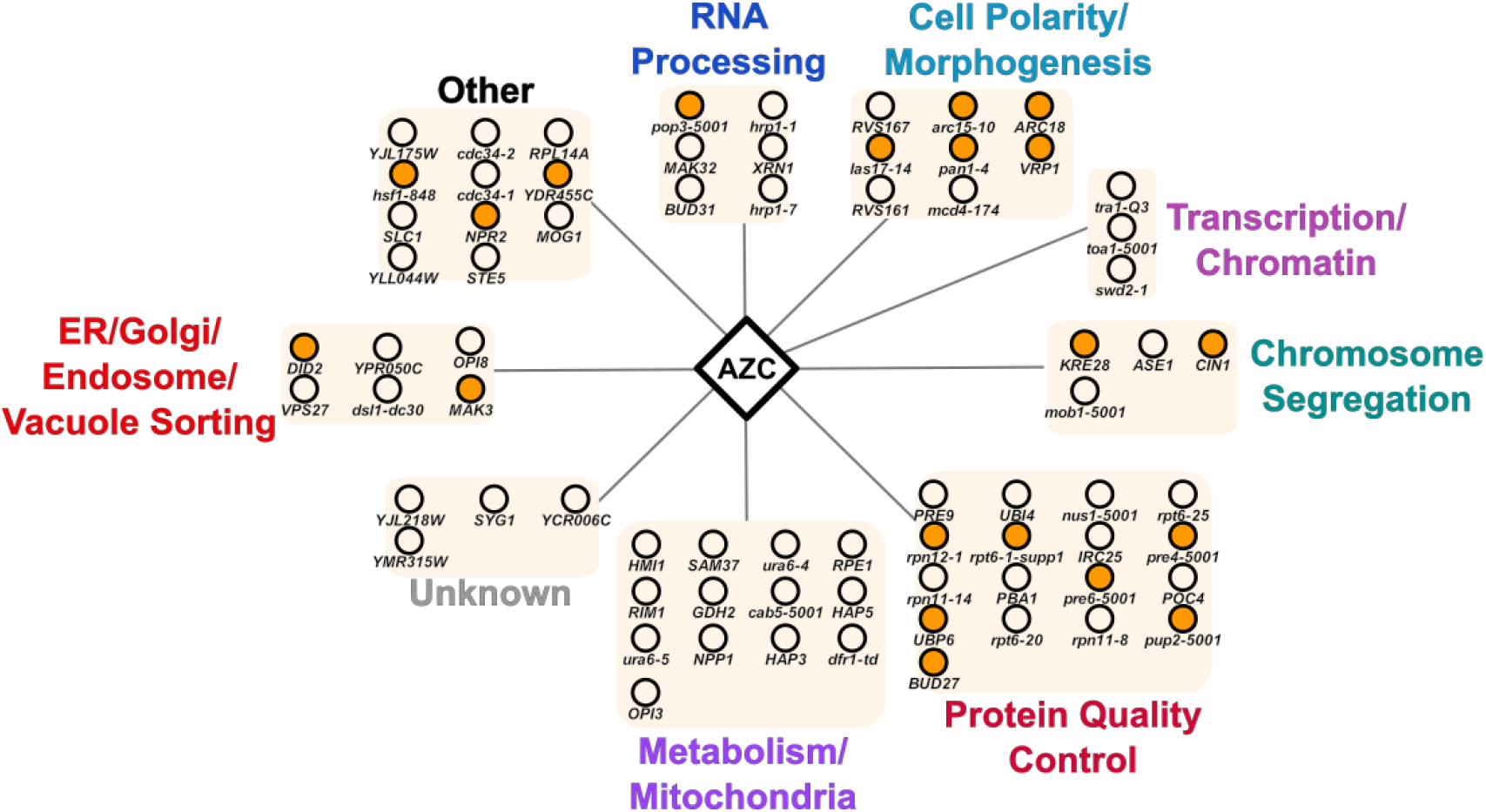
Negative interactions of AZC sensitive genes with tRNA derived serine at proline mistranslation. Alleles that were identified as having negative synthetic interactions with AZC were mated with a strain containing either a wild type tRNA^Ser^ or tRNA^Ser^_UGG, G26A_ which mistranslates serine at proline codons. Double mutants were selected using the SGA method described in Tong *et al*. (2001). Negative genetic interactions (orange) were determined by comparing double mutant fitness of the strains with tRNA^Ser^ to the strains with tRNA^Ser^_UGG, G26A_ in biological triplicate using SGAtools (Wagih *et al*. 2013) and a custom R script (supplemental file 1).

### The proteasome is required to cope with proteotoxic stress caused by mis-incorporation

The protein product of 12 of the genes having negative interactions with AZC either form the proteasome or are involved in the assembly of the proteasome (*UBP6, RPT6, RPN11, PUP2, POC4, PRE6, PBA1, RPN11, PRE4, IRC25, RPN12, PRE9*). This supports previous reports demonstrating the requirement for proteasomal protein degradation for cells to cope with proteotoxic stress (Ruan *et al*. 2008; Hoffman *et al*. 2017; Shcherbakov *et al*. 2019). Consistent with this, the gene encoding ubiquitin (*UBI4*), a post-translational modification required in protein degradation signaling, also has a negative synthetic interaction with AZC and thialysine. These genes as well as *HSF1*, encoding the heat shock response transcription factor, almost certainly are required for cells to cope with proteotoxic stress. We note that no single gene encoding a heat shock chaperone was found to have a negative synthetic interaction with AZC, likely due to genetic redundancy.

### The actin cytoskeleton has a role in protein quality control

In addition to genes involved in protein quality control, our screen identified other genes and pathways either required for resistance to mistranslation or particularly sensitive to mis-made protein. One of these additional pathways was actin patch organization and endocytosis. Interestingly, deletions or temperature sensitive alleles of genes involved in these pathways were sensitive to all three amino acid analogs and tRNA derived mistranslation. These include two genes whose proteins are part of the ARP2/3 complex (*ARC15* and *ARC18*), which is required for motility and integrity of cortical actin patches (Winter *et al*. 1997) and five genes that regulate actin nucleation and endocytosis (*LAS17, VRP1, RVS161, RVS167* and *PAN1;* Naqvi *et al*. 1998; Duncan *et al*. 2001; Sun *et al*. 2006; Youn *et al*. 2010). To determine if AZC alters cellular actin, we imaged cells stained with rhodamine-labelled phalloidin to visualize actin after treatment with various AZC concentrations (Figure 5). The increase in number of actin patches per cell correlated directly with AZC concentration. In yeast, cortical actin patches play roles in endocytosis and cell wall remodelling. Negative synthetic interactions were also observed between amino acid mis-incorporation and *vps27Δ* and *did2Δ*, two genes involved in vacuolar sorting (Piper *et al*. 1995; Nickerson *et al*. 2006). It is possible actin regulation and endocytosis pathways are required to turnover mis-made proteins at the plasma membrane. In support of this, Zhao *et al*. (2013) demonstrated that yeast cells defective in endocytic targeting and exposed to heat shock accumulate mis-folded cell surface protein and die due to loss of plasma membrane integrity. Amino acid mis-incorporation could have a similar effect on cell surface proteins.

**Figure 5.**
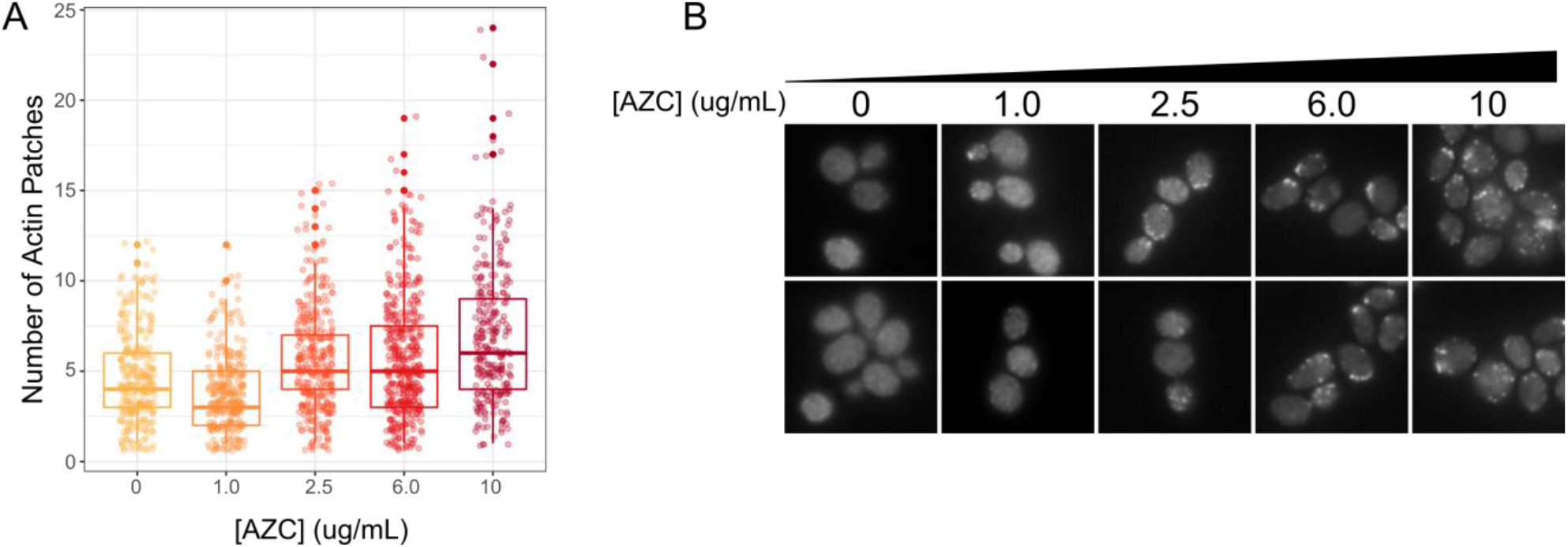
AZC treatment results in increased actin patches. (A) Cells were grown in various concentrations of AZC to an OD_600_ of 0.1, fixed, stained with fluorescein isothiocyanate-ConA, rhodamine-phalloidin, and 4’,6-diamidino-2-phenylindole as described in Ohya *et al*. (2005) and imaged with 100x magnification on a Zeiss Axio Imager Z1 Fluorescent microscope. Number of actin patches per cell was quantified using CalMorph (Ohya *et al*. 2005). In each condition, at least 270 cells were quantified. (B) Representative images of cells quantified in (A).

### Genes involved in response to nitrogen compounds are required for AZC-induced toxicity

The 12 validated strains where AZC-induced toxicity was suppressed corresponded to deletion or mutation of 11 genes (Figure 6A; Supplemental File 3). GO analysis identified enrichment of genes involved in the cellular response to nitrogen compounds as well as TOR signaling (Figure 6B).

**Figure 6.**
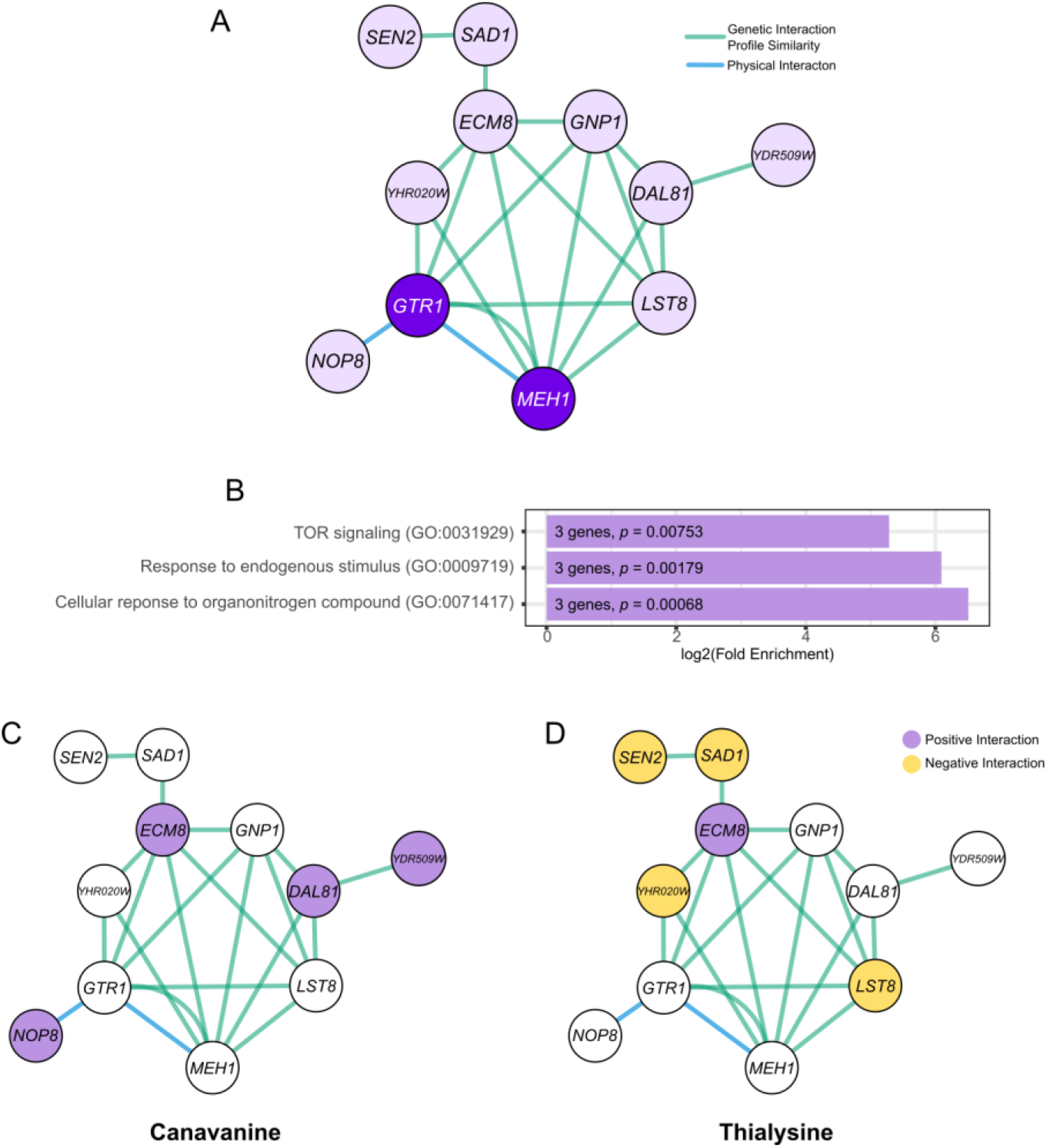
Genes that suppress AZC-induced toxicity. (A) Network of genes required for AZC-induced toxicity. Genetic and physical interaction networks were generated using GeneMANIA. Nodes represent gene/proteins, green edges represent genes that have similar genetic interaction profiles (Costanzo *et al*. 2010, 2016) and blue edges represent proteins that physically interact. Dark purple indicates genes that are part of the EGO complex. (B) Significantly enriched GO biological processes were determined from the set of genes that suppress AZC toxicity. The number of genes corresponding to each GO term and P-values are labelled on each bar. (C) Chemical genetic interaction of genes that suppressed AZC toxicity with canavanine were determined by growth in liquid media. Strains that suppressed canavanine toxicity are colored green. (D) Chemical genetic interaction of genes that suppressed AZC toxicity with thialysine were determined by growth in liquid media. Strains that suppressed thialysine toxicity are colored purple while strains that had negative synthetic interactions with thialysine are colored yellow.

Three of the genes that suppressed AZC toxicity function in AZC import. *GNP1*, a high affinity glutamine permease (Zhu *et al*. 1996), and *YDR509W*, a gene which overlaps with the 5’ coding sequence of *GNP1*, were both identified as suppressors of AZC-induced toxicity. Andréasson *et al*. (2004) demonstrated that Gnp1 is one of the permeases that imports AZC. In addition, AZC toxicity was suppressed by deleting *DAL81. DAL81* is required for *GNP1* expression, through the *S*sy1-*P*tr3-*S*sy5 (SPS) sensing pathway of extracellular amino acids (Boban and Ljungdahl 2007).

We tested if the twelve AZC suppressing alleles more generally suppress mis-incorporation using thialysine and canavanine (Figure 6C and 6D). Only deleting *ECM8* suppressed toxicity resulting from all three non-proteinogenic amino acids. *ECM8* is an uncharacterized gene whose deletion results in resistance to the aminoglycoside hygromycin b, an inhibitor of translation (Lussier *et al*. 1997). It is possible that deleting *ECM8* suppresses mis-made proteins by impacting translation levels. Interestingly, deletions of *YDR509W* and *DAL81*, described above to be involved in AZC uptake, also suppressed canavanine sensitivity suggesting these genes might have a broader role in the uptake of amino acid analogs.

Suppression by mutation of genes involved in the TOR pathway is more specific since *GTR1, MEH1* and *LST8* only suppressed AZC toxicity. Both Gtr1 and Meh1 are part of the EGO complex which regulates sorting of the general amino acid permease Gap1 between the endosome and plasma membrane (Gao and Kaiser 2006). Gtr1 also activates TORC1 in response to amino acid stimulation (Panchaud *et al*. 2013). Downstream of TOR activation, Lst8 inhibits transcription factors that activate enzymes responsible for glutamate and glutamine biosynthesis (Chen and Kaiser 2003). Decreasing either TOR activation or downstream Lst8 activation increases glutamate and glutamine synthesis, which triggers targeting of a variety of amino acid permeases to the vacuole. Roberg *et al*. (1997) found that cells with the *lst8-1* mutation are resistant to AZC, but sensitive to thialysine. Similarly, we observed that both *lst8-6* and *lst8-15* had negative synthetic interactions with thialysine, supporting the idea that Lst8 regulates a specific set of amino acid permeases.

The temperature sensitive allele of *YOR020W*, the prolyl-tRNA synthetase, suppressed only AZC toxicity and likely decreases the amount of AZC charged onto tRNAs. We observed a negative synthetic interaction between this allele and thialysine, likely reflecting the synthetic toxicity of decreasing charged tRNA^Pro^ and mis-incorporating thialysine at lysine codons. It is less clear how the other genes suppress AZC-induced toxicity. It should be noted two strains, *nop8-101* and *sad1-1*, had decreased fitness and that AZC could be suppressing the slow growth caused by the mutation, rather than the mutation suppressing AZC toxicity. Sen2 and Sad1, which only had as positive interaction score with AZC, are involved in the splicing of tRNAs and mRNA, respectively (Winey and Culbertson 1988; Lygerou *et al*. 1999). Interestingly, we observed negative synthetic interactions of *sen2-1* and *sad1-1* with thialysine, indicating that the interaction between these alleles and AZC is not the same for all types of proteotoxic stress. Nop8 is a nucleolar protein required for 60S ribosomal subunit biogenesis and physically interacts with Gtr1 (Zanchin and Goldfarb 1999; Todaka *et al*. 2005). The temperature sensitive *nop8-101* allele also had a positive interaction with canavanine. Nop8 may regulate Gtr1 and act through the TOR pathway discussed above for AZC or may play a more general role in controlling amino acid uptake.

### Conclusions

Using a chemical-genetic analysis we identified genes with positive and negative genetic interactions with AZC. Many of the genes with negative interactions have roles in protein quality control and are involved in the proteasome, suggesting this is the main way cells degrade mistranslated proteins. We also identified genes involved in actin organization, endocytosis and vacuole sorting suggesting a possible role for the recycling of mis-made plasma membrane proteins in preventing toxicity due to mis-incorporation. These same processes are also required for cells to cope with two other non-proteinogenic amino acids, canavanine and thialysine and serine at proline mistranslation caused by a tRNA variant.

Mistranslated proteins, either with non-proteinogenic amino acids through genetic code expansion applications or with canonical amino acids using tRNA and aminoacyl-tRNA synthetase variants, have applications in synthetic biology (Bacher *et al*. 2005; Wang and Pan 2015; Reynolds *et al*. 2017). Incorporating non-proteinogenic amino acids with novel side chains can expand functionality or specificity of a protein. Mistranslation with canonical amino acids yields “statistical proteins”, which are pools of molecules made from the same genetic template but with slightly different amino acid compositions and as such can broaden the function of a protein. Both situations result in proteotoxic stress by globally producing mis-made proteins that must be managed by the cell. Our screen has identified potential genes and pathways that carry out this function. Through modulating these genes and pathways, it may be possible to create strains more tolerate to mis-made protein. Most promising of these are genes with negative genetic interactions because many of those with positive interactions are either specific for AZC and/or would impact translation.

## ACKNOWLEDGMENTS

We would like to thank Patrick Lajoie for use of his plate reader, and the Biotron Integrated Microscopy facility at Western University for their instruction and for use of their fluorescence microscope and Drs. Ohya and Ohnuki for their support with CalMorph software.

